# demuxmix: Demultiplexing oligonucleotide-barcoded single-cell RNA sequencing data with regression mixture models

**DOI:** 10.1101/2023.01.27.525961

**Authors:** Hans-Ulrich Klein

## Abstract

**Motivation:** Droplet-based single-cell RNA sequencing (scRNA-seq) is widely used in biomedical research to interrogate the transcriptomes of single cells on a large scale. Pooling and processing cells from different samples together can reduce costs and batch effects. In order to pool cells, cells are often first labeled with hashtag oligonucleotides (HTOs). These HTOs are sequenced along with the cells’ RNA in the droplets and are subsequently used to computationally assign each droplet to its sample of origin, which is referred to as demultiplexing. Accurate demultiplexing is crucial and can be challenging due to background HTOs, low-quality cells/cell debris, and multiplets.

**Results:** A new demultiplexing method, demuxmix, based on negative binomial regression mixture models is introduced. The method implements two significant improvements. First, demuxmix’s probabilistic classification framework provides error probabilities for droplet assignments that can be used to discard uncertain droplets and inform about the quality of the HTO data and the demultiplexing success. Second, demuxmix utilizes the positive association between detected genes in the RNA library and HTO counts to explain parts of the variance in the HTO data resulting in improved droplet assignments. The improved performance of demuxmix compared to existing demultiplexing methods is assessed on real and simulated data. Finally, the feasibility of accurately demultiplexing experimental designs where non-labeled cells are pooled with labeled cells is demonstrated.

**Availability:** R/Bioconductor package demuxmix (https://doi.org/doi:10.18129/B9.bioc.demuxmix)

## 1 Introduction

Single-cell transcriptomic profiling is fundamental to studying complex biological systems. New droplet-based techniques enable the parallel processing of thousands of cells in a single run (Klein, et al., 2015; Macosko, et al., 2015; Zheng, et al., 2017). Pooling cells from different samples before droplet-based single-cell RNA sequencing (scRNA-seq) can reduce costs and batch effects (Cheng, et al., 2021). If genetically unrelated samples are pooled, common genetic variants can be used to assign droplets to the respective donor after sequencing (Huang, et al., 2019; Kang, et al., 2018). Alternatively, if genetically related or identical samples are pooled, cells can be labeled by hashtag oligonucleotides (HTOs) using oligonucleotide-conjugated antibodies (Stoeckius, et al., 2018) or chemical labeling (Gehring, et al., 2020; McGinnis, et al., 2019). The HTOs are sequenced together with the RNA in the droplet and are subsequently used to assign each droplet to its sample of origin, which is termed demultiplexing.

Demultiplexing is complicated by the presence of multiplets, i.e., droplets with more than one cell. If a multiplet contains cells from different samples, it can be detected based on its HTO profile during demultiplexing. Further, depending on the sample quality, a substantial amount of background HTOs can be observed in some experiments. Background HTOs are HTOs that were not used to tag the cell in the considered droplet. Several methods have been proposed for HTO-based demultiplexing. HTODemux implemented in the R package Seurat clusters cells into groups of low and high counts for each HTO, models the distribution of the cluster with low counts, and then classifies outliers above the 0.99 quantile as tagged (positive) cells (Stoeckius, et al., 2018). hashedDrops in the R/Biocondutor package DropletUtils uses thresholds based on the ratio of the most abundant HTO to the second most abundant HTO to assign droplets to samples. MULTI-seq detects two local maxima in a kernel density estimation for each HTO and defines a threshold between the two maxima such that the number of singlets passing only one of the thresholds across all HTOs is maximized (McGinnis, et al., 2019). GMM-Demux applies a Gaussian mixture model to normalized HTO counts to classify droplets (Xin, et al., 2020). DemuxEM differs from the other methods by explicitly modeling the distribution of background HTO counts using empty droplets (Gaublomme, et al., 2019). All these methods can identify multiplets and, if default parameters are used, may reject to classify some droplets in case of high classification uncertainty.

The new method, demuxmix, described here uses two-component mixture models like GMM-Demux but differs substantially in two important aspects. First, instead of transforming HTO counts to approximate a normal distribution, the negative binomial distribution is used to model raw HTO counts. A good model fit depends on the selected probability distributions and is crucial for accurate class probabilities. Second, demuxmix leverages the association between detected genes and HTO counts in the droplets using regression mixture models to explain some of the variance observed in HTO counts, thereby improving the demultiplexing results. demuxmix is described in detail in the Methods section, and its performance is assessed and compared to the existing methods in the Results section using simulated and real data.

## 2 Methods

For each droplet *i* = 1, …, *m*, a vector *y*_*i,·*_ ∈ ℕ^*n*^ of the counts from the *n* HTOs used in the experiment is observed. Let (*y*_*i,j*_) ∈ ℕ^*m*×*n*^ denote the matrix of HTO counts from all *m* droplets in the experiment. The vector with the numbers of genes detected in the droplets is denoted by *x* ∈ ℕ^*m*^. Any gene with at least one mapped read from the RNA library is considered detected. The task of demultiplexing is to infer the unobserved classes *C*_*i*_ ∈ (0,1}^*n*^ where the *j*-th entry is 1 if droplet *i* contains a cell tagged with the *j*-th HTO. If all entries of *C*_*i*_ are 0, the droplet is empty. If exactly one entry is 1, the droplet is a single-sample droplet (SSD). It can still contain more than one cell from the same sample, which is difficult to detect based on HTO data and not modeled here. If more than one entry is 1, the droplet is a multi-sample droplet (MSD).

### 2.1 demuxmix algorithm

demuxmix consists of three steps: preprocessing, mixture model fitting, and classification. The first two steps are executed independently for each of the *j* = 1, …, *n* HTOs. Empty droplets and low-quality cells should be identified based on their transcriptomic profile and removed before running demuxmix (Lun, et al., 2019).

#### 2.1.1 Preprocessing

*k*-means clustering with *k=2* is applied to the counts *y*_*·,j*_ from the *j*-th HTO after log-transformation to obtain initial class labels. The cluster with the larger mean HTO count is assumed to contain droplets with cells from sample tagged by HTO *j* mainly and is referred to as positive cluster. Droplets in the positive cluster with *y*_*i,j*_ larger than *1*.*5* IQR + third quartile are flagged as outliers. Similarly, droplets from both clusters with *x*_*i*_ more than *1*.*5* IQR above the third quartile are flagged. Outliers are not removed and are still classified by demuxmix but not used for subsequent model fitting.

#### 2.1.2 Regression mixture model

For each HTO *j*, a negative binomial regression mixture model with two components representing negative droplets (not containing cells tagged by HTO *j*) and positive droplets (containing cells tagged by HTO *j*) is used to model the distribution of *y*_*i,j*_ (Wedel and DeSarbo, 2002). Let *h* denote the probability mass function of the negative binomial distribution. The mixture distribution for the *j*-th HTO can be written as

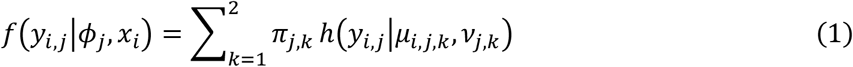

where the vector *ϕ*_*j*_ = (*π*_*j*,1_, *θ*_*j*,1_, *π*_*j*,2_, *θ*_*j*,2_) denotes all model parameters. *π*_*j*,1_, *π*_*j*,2_ > 0, *π*_*j*,1_ + *π*_*j*,2_ = 1, describe the unknown proportions of negative and positive droplets. In a regression mixture model, the means of the observations in each component are predicted from explanatory variables. Here, the number of detected genes in the droplet is used to predict the HTO counts using a negative binomial regression model with canonical link function *g*:

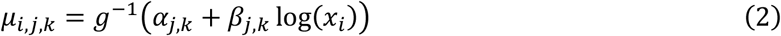

The vector *θ*_*j,k*_ = (*α*_*j,k*_, *β*_*j,k*_, *ν*_*j,k*_) contains the regression coefficients and the dispersion parameter of the *k*-th component.

The model parameters *ϕ*_*j*_ are estimated by the expectation-maximization (EM) algorithm, which iteratively estimates the regression parameters in equation (2) and then updates the class memberships of the droplets. The EM algorithm is initialized using the clustering results from the preprocessing step: droplets from the positive cluster with the larger mean HTO count are assigned to component *k=2*. Thus, component *k=2* represents positive droplets. After the parameters have been estimated, the posterior probability that droplet *i* contains a cell from the sample tagged with HTO *j* can be calculated:

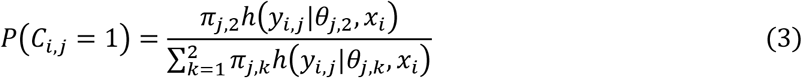

If droplet *i* is more likely to contain a cell from sample *j* than not, then estimated class *ĉ*_*i,j*_ is set to 1 and *ĉ*_*i,j*_ ≔ 0 otherwise.

As shown in the Results section, many datasets demonstrate a positive association between *x* and *y*_*·,j*_, which can be leveraged to improve the demultiplexing results. However, the association can be absent in the negative component if the HTO counts are very low. Moreover, if different cell types with different RNA contents and cell surface properties are pooled, the regression can be driven by the differences between these heterogeneous cell clusters rather than by the association between *x* and *y*_*·,j*_ within each cluster. Therefore, demuxmix fits two simpler mixture models in addition to model (1) and selects the best model for each HTO. The first model does not include a regression model for the negative component *k=1*, i.e., the mean *µ*_*i,j*,1_ in equation (1) becomes *µ*_*j*,1_ and does not depend on the number of detected genes. The second model removes the regression models from both components and is referred to as naïve mixture model. For each HTO, the model which minimizes the expected classification errors calculated using equation (3)

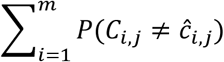

is selected and used for the subsequent droplet classification.

#### 2.1.3 Probabilistic classification

The final classification of a droplet is obtained from the classifications of the *n* separate mixture models *ĉ*_*i*_ = (*ĉ*_*i*,1_, …, *ĉ*_*i,n*_). An advantage of mixture models for classification is that the posterior probabilities for each class can be calculated using equation (3). Assuming that the classification results from the different HTOs are independent, the probability of droplet *i* being from class *ĉ*_*i*_ is

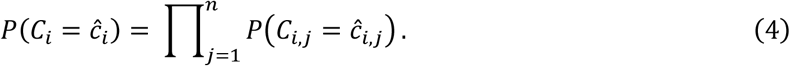

Although the independence assumption is a simplification of the actual data structure, the approximate probabilities from equation (4) are helpful during demultiplexing. In most settings, a droplet with a high classification uncertainty should be discarded and not assigned to a perhaps incorrect sample, which can then negatively affect downstream analyses. demuxmix uses a default threshold of *p*_*acpt*_ = 0.9^*n*^ and classifies droplets as ‘uncertain’ if *P*(*C*_*i*_ = *ĉ*_*i*_) < *p*_*acpt*_. The threshold can be adapted to account for different acceptable demultiplexing error rates in different experimental settings.

### 2.2 Datasets

Two publically available datasets and one new dataset were used to demonstrate demuxmix’s usability. All datasets were generated by the droplet-based 10X Genomics Chromium scRNA-seq platform (Zheng, et al., 2017) using oligonucleotide-labeled antibodies to tag cells before pooling (Stoeckius, et al., 2018).

#### 2.2.1 Human brain single-nucleus dataset

The human brain single-nucleus RNA sequencing (snRNA-seq) dataset consists of nuclei from the dorsolateral prefrontal cortices from 8 genetically unrelated donors. Each sample was tagged with a different HTO. The dataset has been described previously (Gaublomme, et al., 2019). demuxlet was used as an alternative demultiplexing approach based on genetic variation and used as ground truth in the benchmark (Kang, et al., 2018). demuxlet was applied as in the original work: Droplets with a doublet probability > 0.99 were classified as doublets. The remaining droplets were assigned to the most likely singlet class unless they were flagged as ambiguous by demuxlet. The preprocessed dataset of 2,509 high-quality nuclei and the demuxlet results were obtained from the Docker image provided by Gaublomme, et al. (2019).

#### 2.2.2 Cell line mixture dataset

The cell line mixture dataset consists of 12 samples from 4 different cell lines. Each sample was tagged with a different HTO as described previously (Stoeckius, et al., 2018). Droplets were demultiplexed based on the distinct transcription profiles of the 4 cell lines independent of the HTOs using a standard single-cell clustering workflow as described by Amezquita, et al. (2020): Empty droplets were removed using the emptyDrops method (Lun, et al., 2019). The remaining 7,596 droplets were normalized and log-transformed using the *logNormCounts* method from the R package scater. The top 5,000 genes with the largest biological component of variance were selected using *modelGeneVar*. Principal component analysis (PCA) was then applied to the top 500 most variable genes in this set. Droplets were clustered using the Walktrap community detection algorithm applied to the n=10 nearest-neighbor graph constructed from the top 50 principle components (Pons and Latapy, 2006). A total of 19 clusters were detected. The larger clusters (≥ 200 droplets) reflected the different cell lines, and droplets from these clusters were labeled accordingly, except for one cluster of 294 cells, which demonstrated a high amount of mitochondrial reads (mean of 16.6%), indicating apoptotic cells. This cluster and the remaining smaller clusters, some of which likely reflect multiplets, were labeled as ‘uncertain’. The dataset was downloaded from Gene Expression Omnibus (GSM3501446 and GSM3501447).

#### 2.2.3 CSF dataset

The cerebrospinal fluid (CSF) dataset consists of CSF cells pooled with peripheral blood mononuclear cells (PBMCs). Only the PBMCs but not the CSF cells were stained using oligonucleotide-labeled antibodies (BioLegend, 394613) following the manufacturer’s protocol. Cell capture, amplification, and library construction on the 10x Genomics Chromium platform (v3 Chemistry) was performed according to the 10x protocol. Libraries were sequenced on the Illumina NovaSeq instrument and aligned and preprocessed using the Cell Ranger software, including the removal of empty and low-quality droplets. A total of 2,510 droplets with ≥ 200 detected genes were obtained. To allow for genetic demultiplexing using the freemuxlet software, CSF and PBMCs were obtained from two genetically unrelated donors. A droplet was assigned to the respective class ‘CSF’ or ‘PBMC’ if freemuxlet achieved a posterior probability ≥0.99 and labeled ‘uncertain’ otherwise. The study was approved by an Institutional Review Board of Columbia University Irving Medical Center. The dataset is included in the R/Bioconductor package demuxmix.

### 2.3 Simulation

Simulated data to benchmark the demultiplexing methods were generated based on real datasets using only high-confidence SSDs as starting set. For the human brain dataset, high-confidence SSDs were defined as droplets identified as SSDs by the genetic demultiplexing (demuxlet) with a droplet probability < 0.1 and a probability of the most-likely singlet class > 0.999. This stringent filter resulted in 2,123 SSDs with known sample source. To simulate low-quality datasets, a scaling factor *s* = 1, 0.95, …, 0.1 was used to attenuate the true HTO signal: 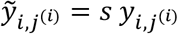 where *j*^(*i*)^ refers to the HTO used to tag the cell in droplet *i*. Background HTO counts were not altered. After attenuating the HTO signal, 200 doublets were generated by randomly merging two SSDs from different donors. The HTO and RNA read counts of the artificial doublets were set to the average values of the two merged SSDs.

Similarly, data were simulated based on the cell line mixture dataset. All 7,047 droplets assigned to one of the 4 cell lines based on their RNA profile as described in section 2.2.2 were considered high-confidence SSDs and used for the simulation. The scaling factor ranged from 1 to 0.25 since all methods performed poorly on this dataset with *s* < 0.25. The clustering-based ground truth only identified the cell line but not which of the three HTOs used for each cell line was used to tag the cell in the droplet. Thus, to decide which HTO should be multiplied by the scaling factor *s*, majority voting was applied after running all 6 methods included in the benchmark on the unmodified dataset. In case of a tie (observed for 66 out of 7,047 droplets), one of the top-voted HTOs was randomly drawn. A total of 350 doublets were generated and added to the dataset in the same way as in the human brain dataset.

## 3 Results

Three different HTO datasets from three laboratories were used to assess demuxmix’s performance. The first dataset is the human brain single-nucleus dataset, which consists of pooled nuclei isolated from 8 different donors. A different HTO was used for each donor (Fig. 1A). The median number of HTO reads per droplet was 1,454. Genetic demultiplexing was applied and used as ground truth.

**Fig. 1.**
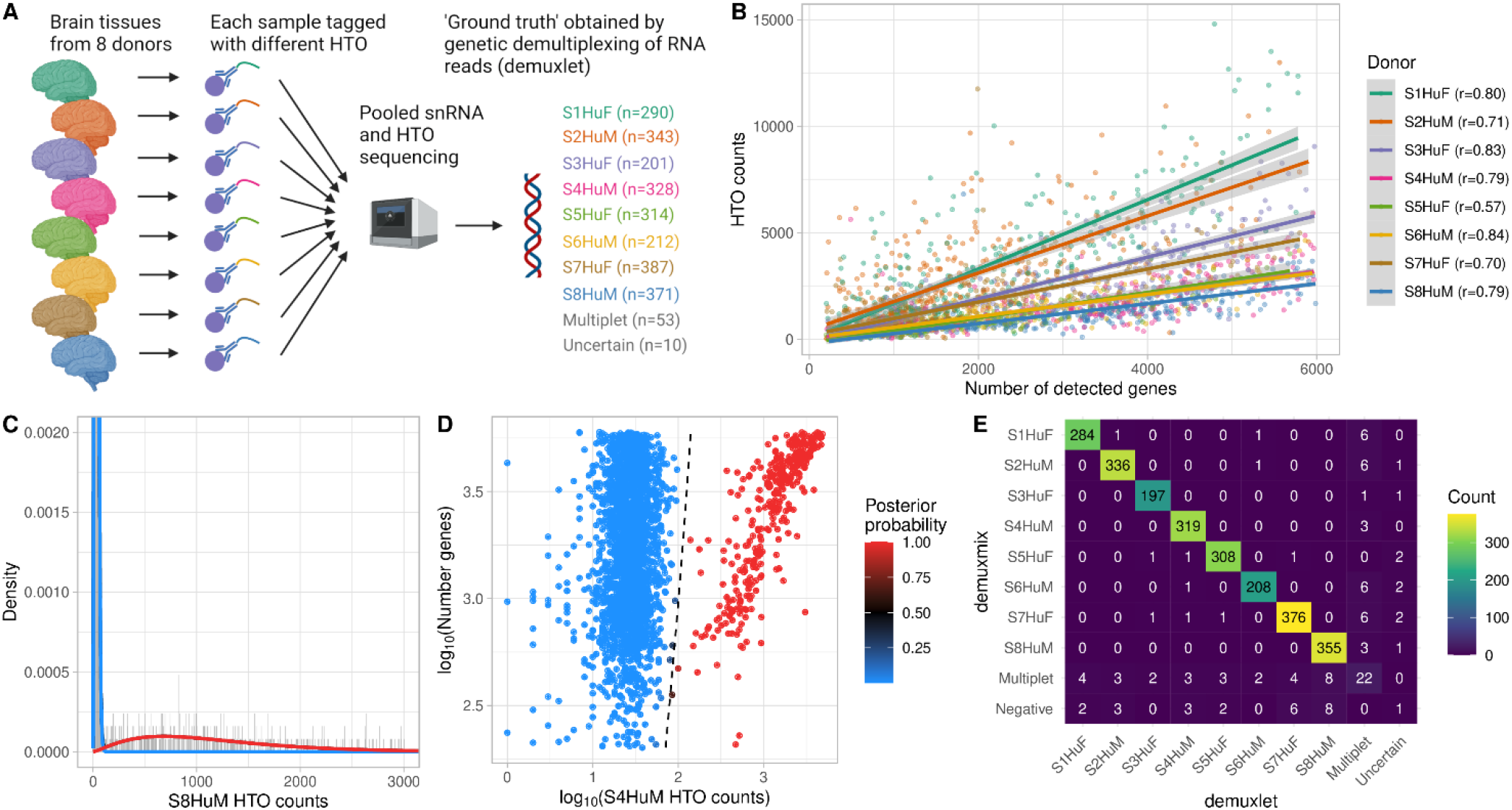
demuxmix results from the human brain single-nucleus dataset. **A)** Experimental design of the human brain single-nucleus dataset. **B)** Scatter plot shows the correlation between the number of detected genes and HTO counts. Shown are the counts of the HTO used to tag the respective nucleus as determined by genetic demultiplexing. Regression lines are plotted, and Pearson correlation coefficients are shown in the legend. **C)** Histogram depicts the distribution of an exemplary HTO (S8HuM) overlaid by the probability mass functions of the negative (blue) and the positive (red) component of the mixture model. **D)** Scatter plot shows the relation between HTO counts (x-axis), the number of detected genes (y-axis), and the posterior probability for the droplet containing a cell tagged by the HTO (color) obtained from the model. The black dashed curve indicates the decision boundary where the posterior probability equals 0.5. **E)** Matrix displays the concordance between the droplet classifications obtained from genetic demultiplexing by demuxlet (x-axis) and demuxmix (y-axis).

### 3.1 HTO counts are positively correlated with the number of detected genes in the droplets

To study the association between the number of detected genes and the HTO count in a droplet, the number of genes was plotted against the number of reads of the HTO used to tag the respective nucleus (Fig 1B). All samples demonstrated a positive correlation ranging from 0.57 to 0.84, indicating that the number of detected genes explains parts of the variance in HTO counts. This association is likely driven by biological factors, e.g., nucleus size, and technical factors like the quality of the gel beads. A positive correlation was also observed for background HTO reads (HTOs not used to tag the respective nucleus), albeit with a much lower correlation ranging from 0.04 to 0.09, likely due to the low background HTOs counts (median of 25 reads).

### 3.2 demuxmix’s classification is highly consistent with genetic demultiplexing

For 6 out of 8 HTOs, demuxmix selected the mixture model with a regression model in the positive but not in the negative component, reflecting the observed weak correlations between background HTO counts and number of detected genes. The naïve mixture model and the mixture model with two regression components were each selected once. Figure 1C exemplarily shows the histogram for one HTO (S8HuM) overlaid with the probability mass functions of the mixture model. The dataset was originally generated to demonstrate the feasibility of tagging single nuclei and is of exceptional quality. The negative component (blue) shows very low background HTO counts and little overlap with the positive (red) component. Figure 1D shows the decision boundary for the HTO S4HuM. Since a regression mixture model was selected for this HTO, the decision boundary is a curve depending on both HTO counts and the number of genes. demuxmix’s classification results were highly concordant with those from genetic demultiplexing (Fig. 1E).

### 3.3 demuxmix demonstrates superior performance on simulated benchmark data

To systematically assess and compare the performance of demuxmix to previously published demultiplexing algorithms, HTO datasets were simulated based on droplets identified by demuxlet as high-confidence SSDs. The HTO signal in the data was then gradually attenuated by a scaling factor *s* to simulate low-quality data. MSMs were artificially generated by merging two SSDs. To assess the methods’ performances, the probability that the predicted class *ĉ*_*i*_ is correct given the method predicted an SSD (*P*(*C*_*i*_ = *ĉ*_*i*_ |*ĉ*_*i*_ ∈ SSD), referred to as precision_SSD_) was calculated. This measure is essential in the context of demultiplexing, because falsely predicted SSDs remain in the dataset and can negatively affect downstream analyses. Since a high precision_SSD_ can be achieved by generously classifying ambiguous droplets as ‘uncertain’ or as MSMs, precision_SSD_ was considered together with the sensitivity_SSD_, which was defined as the probability that a true SSD was identified as SSD, i.e., *P*(*ĉ*_*i*_ ∈ SSD|*C*_*i*_ ∈ SSD). A low sensitivity_SSD_ usually leads to many droplets being removed from the dataset. Supplementary Figures S1A and B show the two performance measures for different scaling factors *s* ranging from 1 to 0.1. Table 1 shows the average measures observed across all values of *s*. All methods were applied using default parameters. Notably, all methods achieved a high precision_SSD_ for larger values of *s*, reflecting the high quality of the dataset. While differences in the average precision_SSD_ were relatively small between methods, larger differences were observed for the average sensitivity_SSD_ (Table 1). demuxmix achieved the highest average precision_SSD_ and the highest average sensitivity_SSD_. Only 6.3% of true SSDs were discarded as ‘uncertain’, ‘negative’, or MSMs by demuxmix, whereas other methods, on average, discarded up to 12.8% of true SSDs.

**Table 1.**
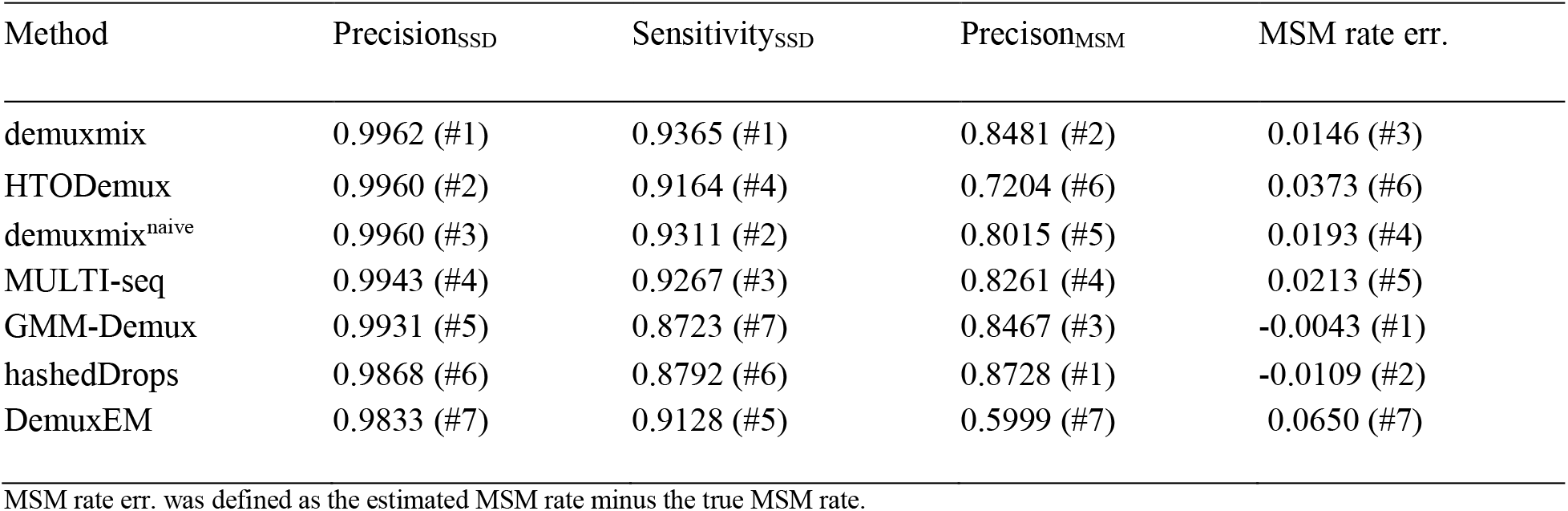
Benchmark results from the human brain dataset

| Method | Precision<sub>SSD</sub> | Sensitivity<sub>SSD</sub> | Precision<sub>MSM</sub> | MSM rate err. |
| --- | --- | --- | --- | --- |
| demuxmix | 0.9962 (#1) | 0.9365 (#1) | 0.8481 (#2) | 0.0146 (#3) |
| HTODemux | 0.9960 (#2) | 0.9164 (#4) | 0.7204 (#6) | 0.0373 (#6) |
| demuxmix<sup>naïve</sup> | 0.9960 (#3) | 0.9311 (#2) | 0.8015 (#5) | 0.0193 (#4) |
| MULTI-seq | 0.9943 (#4) | 0.9267 (#3) | 0.8261 (#4) | 0.0213 (#5) |
| GMM-Demux | 0.9931 (#5) | 0.8723 (#7) | 0.8467 (#3) | -0.0043 (#1) |
| hashedDrops | 0.9868 (#6) | 0.8792 (#6) | 0.8728 (#1) | -0.0109 (#2) |
| DemuxEM | 0.9833 (#7) | 0.9128 (#5) | 0.5999 (#7) | 0.0650 (#7) |

MSM rate err. was defined as the estimated MSM rate minus the true MSM rate.

While high precision and sensitivity for SSDs are most important, identifying MSMs reliably can be helpful for two reasons. First, MSMs can be used to detect single-sample multiplets (SSMs) with similar transcription profiles in downstream clustering analyses. Second, accurately estimating the dataset’s MSM rate can help plan future experiments. Therefore, two additional measures precision_MSM_ and MSM rate were calculated. Precision_MSM_ was defined as the probability that a droplet classified as MSM is a true MSM, i.e., *P*(*C*_*i*_ ∈ MSM|*ĉ*_*i*_ ∈ MSM). The MSM rate was calculated by dividing the number of predicted MSMs by the number of droplets in the dataset after removing droplets flagged as ‘uncertain’ by the respective method. As shown in Table 1, all methods perform worse in classifying MSMs, and a larger fraction of predicted MSMs (13-40%) were actual SSDs. Most methods slightly overestimated the MSM rate. Detailed results are shown in Supplementary Figures S1C and D.

### 3.4 demuxmix performs better than naïve mixture models

A modified version of demuxmix using naïve mixture models exclusively (demuxmix naïve) was run on the benchmark data to investigate whether regression mixture models perform better than naïve mixture models. Although demuxmix naïve still performed better than some of the other methods, it performed worse than demuxmix according to the four measures used in this benchmark, indicating that the use of regression mixture models improved the demultiplexing of this dataset (Table 1, Supplementary Figures S1A-D). The use of regression mixture models improved the detection of MSMs in particular. Further, the better SSD classification of demuxmix naïve compared to GMM-Demux, which uses normal mixture models after data transformation, suggests that the negative binomial mixture models may provide a better data fit.

### 3.5 Validation in independent dataset

The cell line mixture dataset (Stoeckius, et al., 2018) consists of a pool of 12 samples from 4 different cell lines as shown in Figure 2A, and was used to validate demuxmix’s performance. Instead of genetic demultiplexing, the cell lines’ distinct transcription profiles were used to assign droplets to cell lines independent of the HTO data. Small clusters potentially consisting of doublets or apoptotic cells were removed, leaving four large distinct clusters representing the four cell lines (Fig. 2A). The median HTO count in this dataset was 185 reads per droplet.

**Fig. 2.**
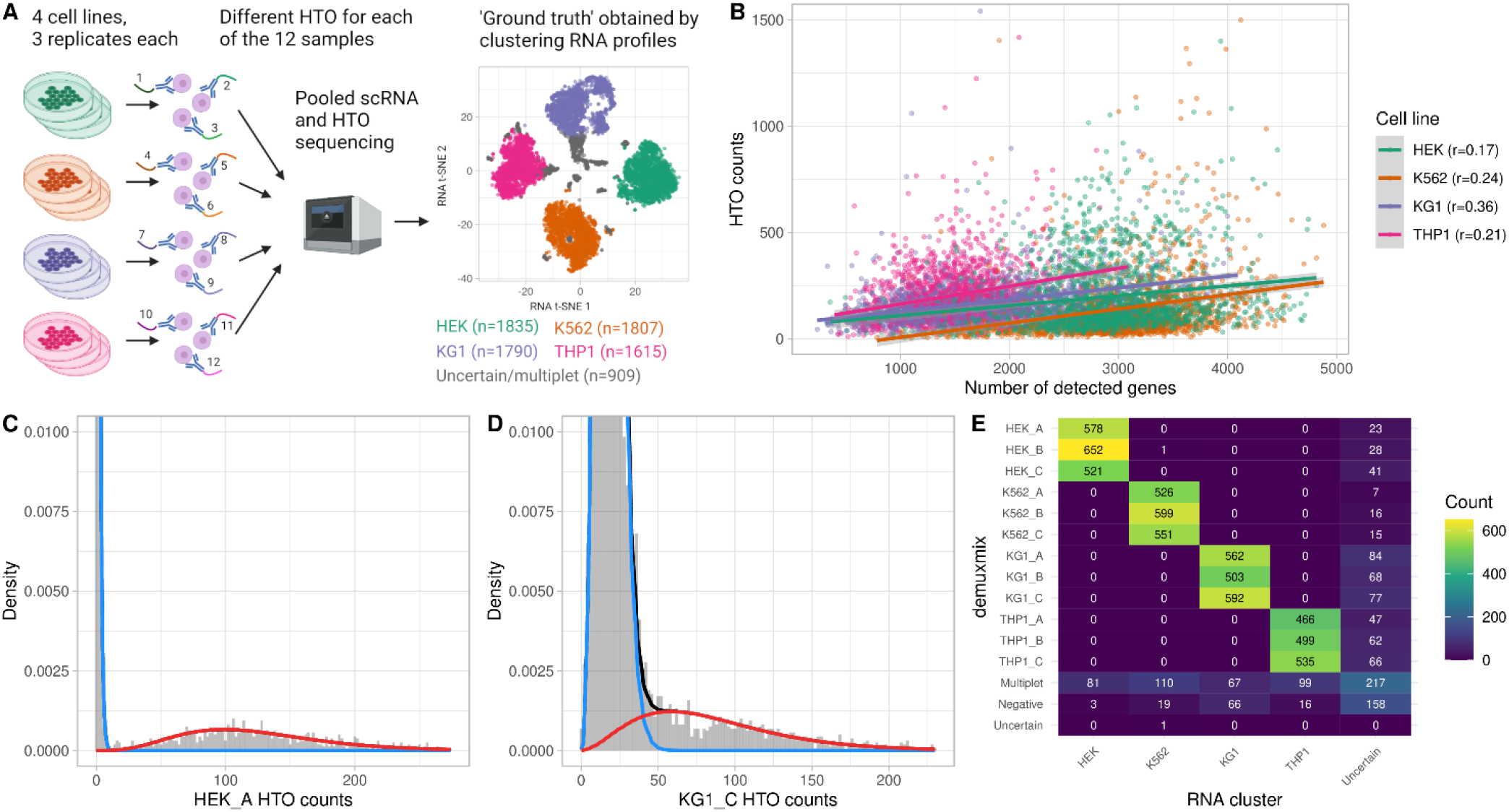
demuxmix results from the cell line mixture dataset. **A)** Experimental design of the cell line mixture dataset. **B)** Scatter plot shows the correlation between the number of detected genes and HTO counts. Shown are the sums of the three HTOs used to tag the respective cell line of origin as determined by transcriptomic clustering. Regression lines are plotted, and Pearson correlation coefficients are shown in the legend. **C)** Histogram depicts the distribution of HTO counts for an exemplary HTO (HEK_A) overlaid by the probability mass functions of the negative (blue) and the positive (red) component of the mixture model. **D)** Histogram as in C) shows the distribution of HTO KG1_C, which demonstrated a larger overlap between the two mixture components. **E)** Matrix displays the concordance between the droplet classifications obtained from transcriptomic clustering (x-axis) and demuxmix (y-axis).

The correlation between HTO counts and the number of detected genes was weaker than in the human brain dataset (Fig. 2B). This was likely caused by the homogeneity of the cells within each cell line, whereas the brain dataset consisted of a mix of neuronal and glial nuclei of different sizes. Thus, the observed weaker correlation is probably mainly driven by technical factors rather than biological differences between the cells. Consequently, demuxmix selected a regression mixture model for only 2 out of the 12 HTOs indicating that modeling the relation between HTO counts and the number of genes improves the demultiplexing only marginally in this dataset. As exemplarily displayed in Figures 2C and D, demuxmix’s models fit the data well. One HTO (‘KG1_C’) showed a larger amount of background HTOs, causing some overlap between the positive and the negative component (Fig. 2D). Still, demuxmix’s classification was highly consistent with the results from clustering the transcriptomic profiles (Fig. 2E). The lower quality of the HTO ‘KG1_C’ (Fig. 2D) resulted in a larger number of KG1 cells classified as ‘negative’ but not in the assignment of incorrect SSD classes (Fig. 2E).

Using the labels from the RNA clustering as ground truth, the cell line mixture dataset was used similarly to the human brain dataset to simulate benchmark data. A total of 350 MSMs were generated, and the HTO signal was attenuated by a scaling factor *s* between 1 and 0.25. The performance of all methods declined considerably for *s* < 0.25. Supplementary Figures S2A-D show the results for each value of *s*; Table 2 shows the average performances. The results mainly replicate the findings from the human brain dataset. demuxmix achieved the highest precision_SSD_ and the highest sensitivity_SSD_. Whereas all methods demonstrate a high precision_SSD_, larger differences were observed in the sensitivity_SSD_, indicating that other methods, compared to demuxmix, discard more SSDs resulting in a loss of data. Similarly, the observed MSM-related metrics were lower than in the human brain dataset, reflecting this dataset’s slightly lower quality and sequencing depth. As expected, the difference between demuxmix and the naïve negative binomial mixture models is small since demuxmix mainly relied on naïve mixture models for this dataset. In summary, these results underline demuxmix’s good performance and its flexibility to adapt to different datasets.

**Table 2.**
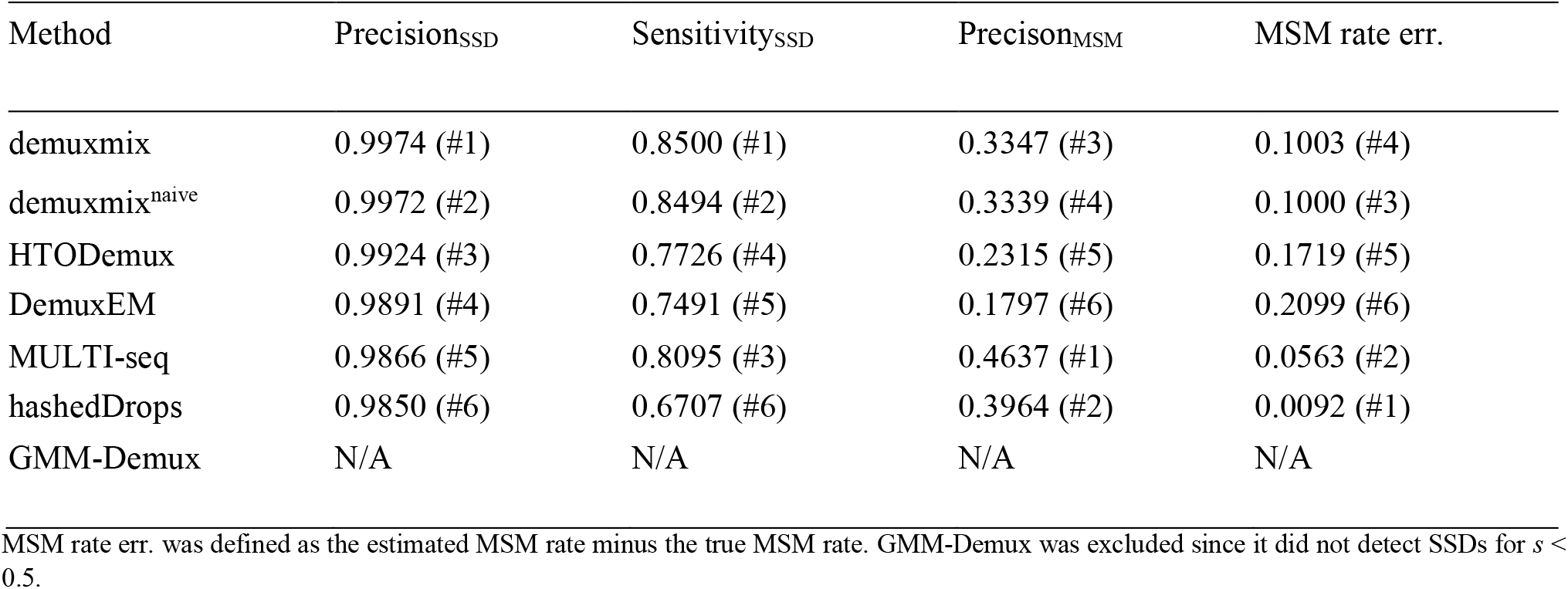
Benchmark results from the cell line mixture dataset

### 3.6 Experimental design with non-tagged cells

Staining cells with oligonucleotide-labeled antibodies requires an additional experimental step, which may alter cell states or lead to cell loss. Thus, when rare and precious cells are pooled with highly abundant cells, staining only the abundant cells is desirable if the demultiplexing algorithm can reliably separate the negative from the positive (tagged) cells. For example, Figure 3A illustrates an experiment where rare untreated CSF cells were pooled with oligonucleotide-labeled PBMCs. To assess demuxmix’s performance in this setting, CSF and blood were obtained from two unrelated donors so that genetic demultiplexing could be used as ground truth.

**Fig. 3.**
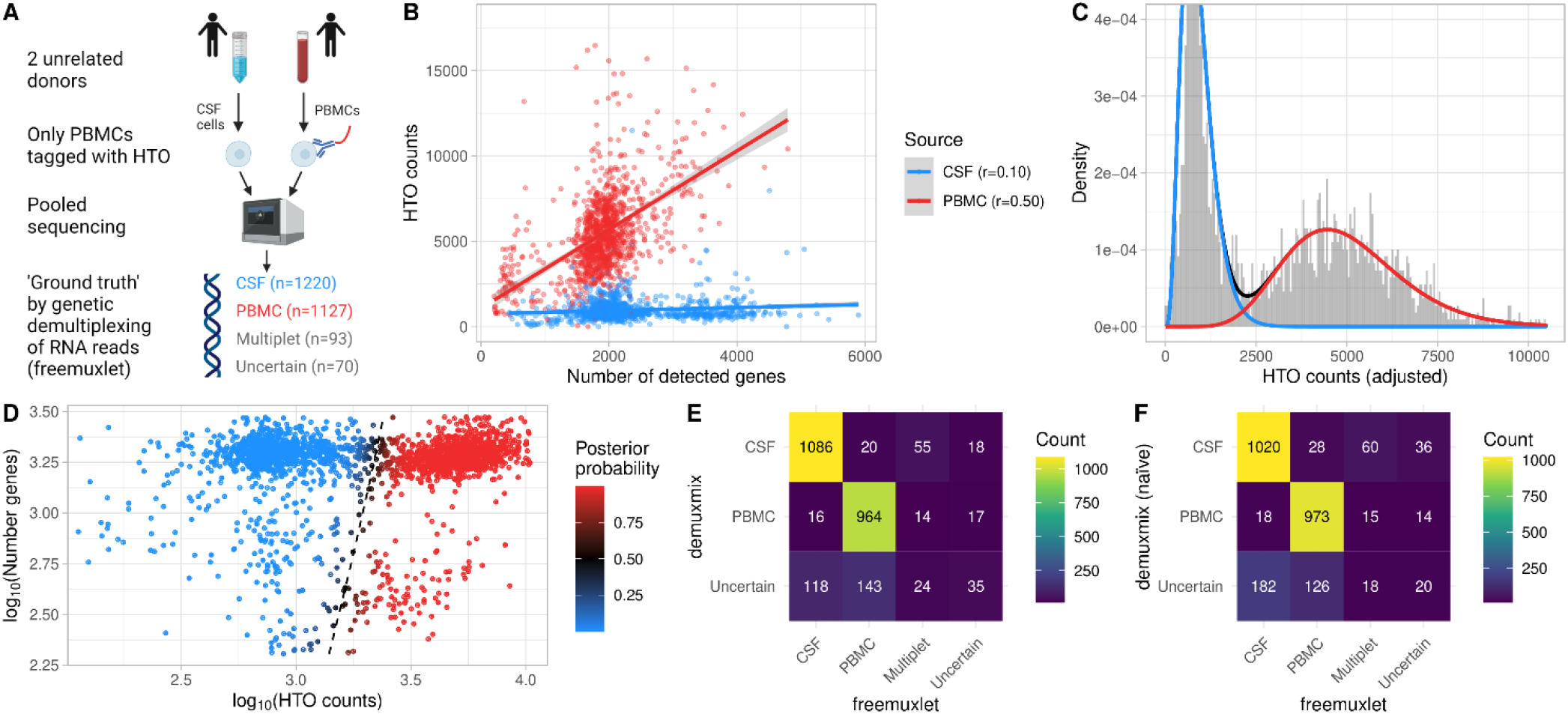
demuxmix results from the CSF dataset. **A)** Experimental design of the CSF dataset. **B)** Scatter plot shows the correlation between the number of detected genes and HTO counts in the droplets. Each droplet is colored by the cell’s source as determined by genetic demultiplexing. Multiplets and uncertain droplets were removed. Regression lines are plotted, and Pearson correlation coefficients are shown in the legend. **C)** Histogram depicts the distribution of HTO counts overlaid by the probability mass functions from the two components of the regression mixture model. Since the mixture probability mass function depends on the number of detected genes, the probability mass function for a droplet with the average number of genes observed in the data is shown, and the histogram shows adjusted HTO counts after regressing out the effect of the number of genes. **D)** Scatter plot shows the relation between HTO counts (x-axis), the number of detected genes (y-axis), and the posterior probability for the droplet containing a PBMC (color) obtained from the model. The black dashed curve indicates the decision boundary where the posterior probability equals 0.5. **E)** Matrix displays the concordance between the droplet classifications obtained from genetic demultiplexing by freemuxlet (x-axis) and demuxmix (y-axis). **F)** As in (E) but only naïve mixture models were used by demuxmix, ignoring the number of detected genes in the droplets.

A positive correlation of 0.50 was observed between the number of genes and HTO counts in the PBMCs (Fig. 3B). In the non-tagged CSF cells, which harbored some background HTOs, the correlation was weaker (*r* = 0.10). In this dataset, demuxmix selected a mixture model with regression models in both components. Figure 3C shows the mixture probability mass function for a droplet with the average number of 1,796.5 detected genes. For such an average droplet, the expected HTO count was 4977.8 for PBMCs and 897.4 for CSF cells. Figure 3D shows the decision boundary of the regression mixture model. For this special design with one non-tagged sample, the acceptance probability was set to *p*_*acpt*_ = 0.99 to increase the precision at the cost of a larger amount of droplets being discarded as ‘uncertain’. Ignoring droplets identified as ‘multiplets’ or ‘uncertain’ by the genetic demultipexing algorithm, demuxmix achieved a specificity of 0.985, a sensitivity of 0.980, and removed 261 cells (11.1%) as ‘uncertain’ (Fig. 3E). In line with the previous observations, results from the naïve mixture models were worse, indicating the benefit of modeling the number of detected genes as explanatory variable (Fig. 3F, specificity of 0.983, sensitivity of 0.972, 308 cells (13.1%) removed as ‘uncertain’). Overall, these results demonstrate the feasibility of this experimental design where sensitive untreated cells are pooled with tagged cells.

## 4 Discussion

Successful demultiplexing is essential for downstream data analyses. The overall good performance of the new method presented here can be attributed to two major improvements: (i) an accurate probabilistic model of HTO counts based on the negative binomial distribution and (ii) the addition of the number of detected features in the RNA library to the model as an explanatory variable that explains parts of the variance of the HTO counts.

The performance of demuxmix was assessed in a comparative study based on two published benchmark datasets with an available ground truth derived from two different approaches utilizing genetic diversity or distinct transcription profiles, respectively. The characteristics of the benchmark datasets have implications for the interpretation of the results. First, the human brain and the cell line mixture datasets were carefully optimized and generated to feature antibody-based nucleus and cell labeling and are of exceptional quality (Gaublomme, et al., 2019; Stoeckius, et al., 2018). Thus, all methods performed well on these data, and the HTO signal had to be attenuated to detect larger differences between the methods. However, typical HTO data is often noisier with more background HTO reads and borderline droplets, as seen in the CSF dataset (Fig. 3D), so the selected method likely has a larger effect on the results. Second, cells in the cell line mixture dataset were distinct between samples but very homogenous within samples (same cell line). This design facilitated demultiplexing based on the transcription profiles but is uncommon in real experimental designs. Cellular heterogeneity is a driving factor underlying the association between HTO counts and the number of detected genes, which could not be utilized by demuxmix in this dataset. Third, the reported error rates in the benchmark study are an upper boundary since the ground truth cannot be assumed to be free of any error despite the stringent filters. Compared to typical experimental designs, the unusually large number of pooled samples (8 and 12) further complicated the demultiplexing. However, these factors are likely offset by the high quality of the benchmark datasets.

The benchmark study provided valuable insights, and despite the differences between the two benchmark datasets, the rankings based on average performance measures were highly similar. Not all demultiplexing errors are equally important. This study primarily focused on (i) whether predicted SSDs were classified correctly (precision_SSD_) since only these droplets remain in the dataset and (ii) on the recovery of SSDs (sensitivity_SSD_) since not detected SSDs result in a loss of data. Notably, the observed precision_SSD_ never fell under 0.9 in the benchmark study, indicating that none of the methods produced many incorrect SSD assignments even when the HTO signal was artificially weakened. However, some methods discarded a significant number of true SSDs as uncertain or multiplets (low sensitivity_SSD_) to achieve a high precision_SSD_. demuxmix demonstrated the highest precision_SSD_ in both datasets and classified on average 94% of SSDs (other methods 87%-93%) in the human brain dataset and 85% of SSDs (other methods 67%-81%) in the cell line mixture dataset.

Finally, the CSF dataset was used to demonstrate that experimental designs where only one of two pooled samples was tagged by HTOs can be successfully demultiplexed by demuxmix. Such a design is useful when pooling rare and sensitive cells together with highly abundant cells.

In summary, demuxmix is a new flexible demultiplexing method that showed superior performance in the benchmark presented here. The method is implemented as an R/Bioconductor package and includes methods for estimating error rates and generating diagnostic plots.

## Acknowledgements

I would like to thank Drs. Victoria S. Marshe and John F. Tuddenham for testing the method on various real datasets and for their valuable feedback, and Dr. Mariko F. Taga for sharing her expertise with cell-hashing experiments. Figures 1A, 2A, and 3A were created with BioRender.com.

