## Supplementary Figures for "demuxmix: Demultiplexing oligonucleotide-barcoded single-cell RNA sequencing data with regression mixture models"

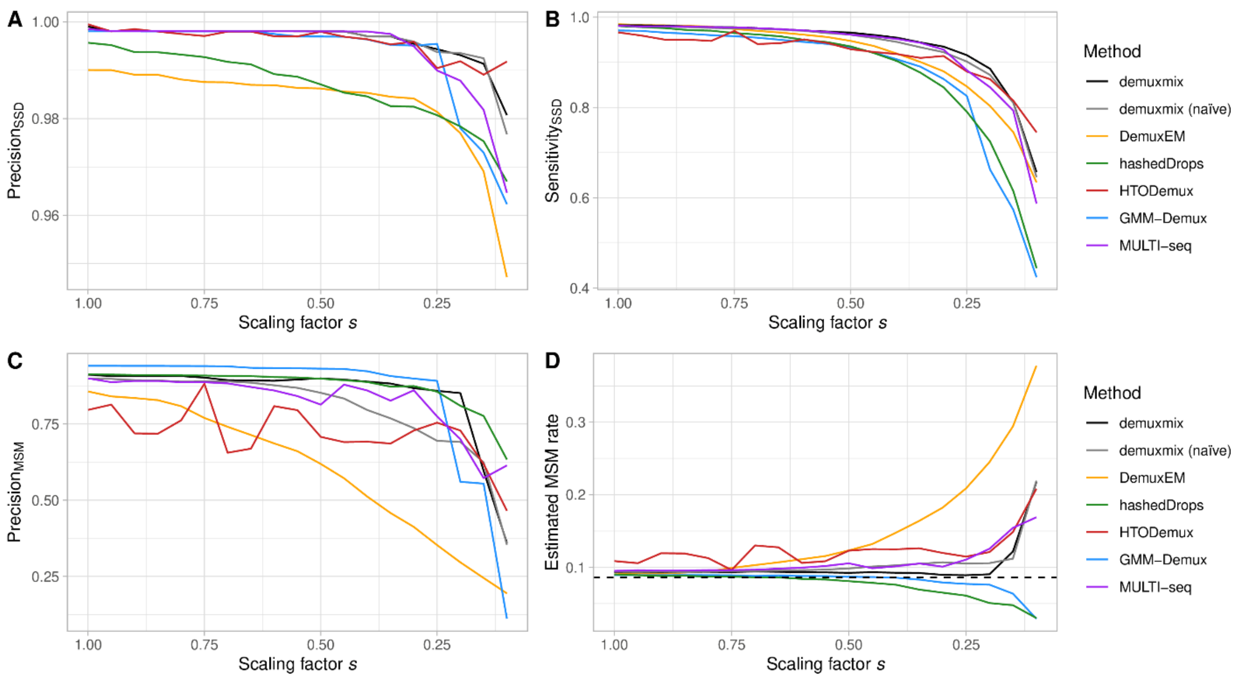


**Supplementary Fig. S1. Benchmark results from the human brain dataset. A)** Plot shows the precision$_{SSD}$ on the y-axis for different scaling factors $s$ on the x-axis. The read counts of the HTO used to tag the cell but not the background HTO counts were multiplied with $s$ to reduce the HTO signal. Precision$_{SSD}$ was defined as the probability that the classification is correct given an SSD class was predicted: $P(C_i = \hat{c}_i | \hat{c}_i \in \text{SSD})$. **B)** Plot shows the sensitivity$_{SSD}$ on the y-axis for different scaling factors $s$ on the x-axis. Sensitivity$_{SSD}$ was defined as the probability that a true SSD was assigned to any SSD class: $P(\hat{c}_i \in \text{SSD} | C_i \in \text{SSD})$. **C)** Plot shows the precision$_{MSM}$ on the y-axis for different scaling factors $s$ on the x-axis. Precision$_{MSM}$ was defined as the probability that a predicted MSM is truly an MSM: $P(C_i \in \text{MSM} | \hat{c}_i \in \text{MSM})$. **D)** Plot shows the estimated MSM rate on the y-axis for different scaling factors $s$ on the x-axis. The true MSM rate in the simulated data is shown as black dashed line.

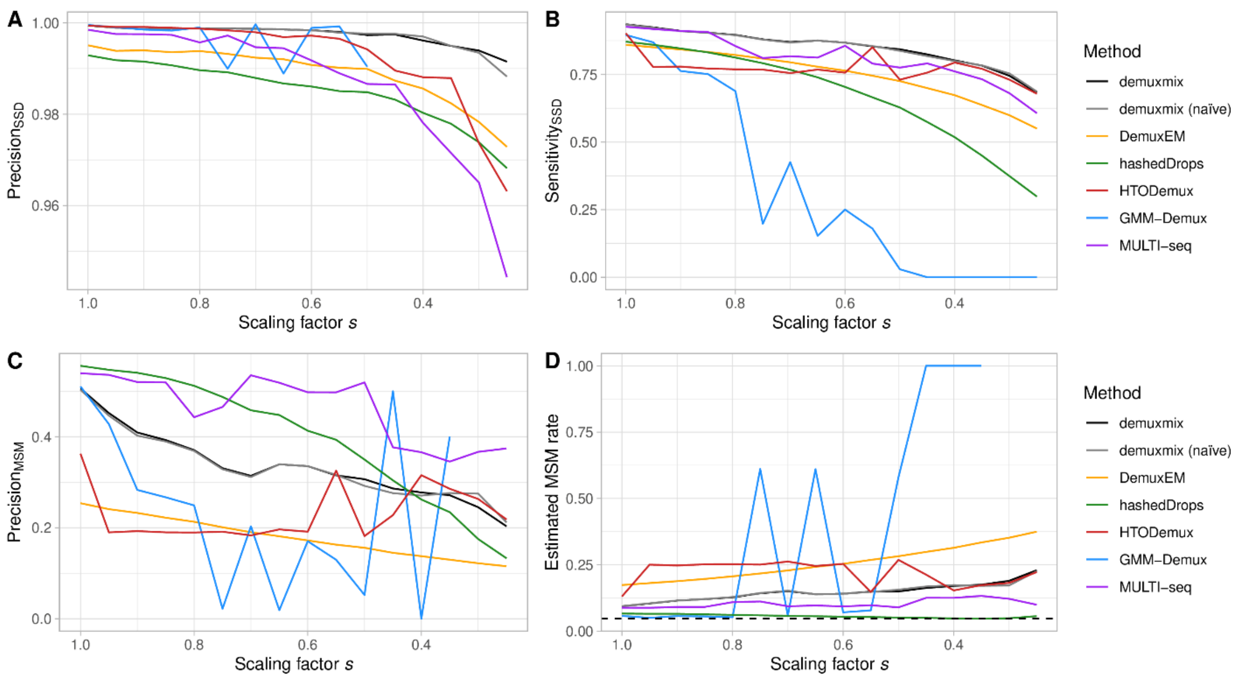


**Supplementary Fig. S2. Benchmark results from the cell line mixture dataset. A)** Plot shows the precision$_{\text{SSD}}$ on the y-axis for different scaling factors *s* on the x-axis. The read counts of the HTO used to tag the cell but not the background HTO counts were multiplied with *s* to reduce the HTO signal. Precision$_{\text{SSD}}$ was defined as the probability that the classification is correct given an SSD class was predicted: $P(C_i = \hat{c}_i | \hat{c}_i \in \text{SSD})$. **B)** Plot shows the sensitivity$_{\text{SSD}}$ on the y-axis for different scaling factors *s* on the x-axis. Sensitivity$_{\text{SSD}}$ was defined as the probability that a true SSD was assigned to any SSD class: $P(\hat{c}_i \in \text{SSD} | C_i \in \text{SSD})$. **C)** Plot shows the precision$_{\text{MSM}}$ on the y-axis for different scaling factors *s* on the x-axis. Precision$_{\text{MSM}}$ was defined as the probability that a predicted MSM is truly an MSM: $P(C_i \in \text{MSM} | \hat{c}_i \in \text{MSM})$. **D)** Plot shows the estimated MSM rate on the y-axis for different scaling factors *s* on the x-axis. The true MSM rate in the simulated data is shown as black dashed line.
